# Overprecision Increases Subsequent Surprise

**DOI:** 10.1101/2019.12.13.875203

**Authors:** Derek Schatz, Don A. Moore

**Affiliations:** University of California at Berkeley

## Abstract

Overconfident people should be surprised that they are so often wrong. Are they? Three studies examined the relationship between confidence and surprise in order to shed light on the psychology of overprecision in judgment. Participants reported ex-ante confidence in their beliefs, and after receiving accuracy feedback, they then reported ex-post surprise. Results show that more ex-ante confidence produces less ex-post surprise for correct answers; this relationship reverses for incorrect answers. However, this sensible pattern only holds for some measures of confidence; it fails for confidence-interval measures. The results can help explain the robust durability of overprecision in judgment.

---

Overprecision is overconfidence in the accuracy of one’s beliefs (Moore, Tenney, & Haran, 2016). This excessive certainty is on display we are too sure we are right (Moore et al., 2017), when people think they know how their friends will behave (Dunning, Griffin, Milojkovic, & Ross, 1990), when doctors are too certain of a favored diagnosis (Arkes, Wortmann, Saville, & Harkness, 1981), or when managers issue excessively precise and inaccurate earnings forecasts (Hribar & Yang, 2015). Frequent feedback should help people calibrate their confidence; being routinely wrong should reduce confidence in the next forecast. However, the robustness and durability of overprecision suggests this corrective may be incomplete (Harvey, 1997; Moore et al., 2016). These overly precise beliefs increase the risk of being wrong. When their expectations prove wrong, people ought to be surprised. Are they? In this paper, we ask how overconfidence contributes to subsequent surprise.

## Overprecision

There are innumerable ways in which overly certain beliefs can impair decisions. Overprecision contributes, for instance, to managers issuing too much debt when they underestimate the volatility of their firm’s future (Hackbarth, 2008), foregoing accounting corrections to their own forecasts of firm returns (Ahmed & Duellman, 2013), and issuing excessively precise and inaccurate earnings forecasts (Hayward & Fitza, 2017; Hribar & Yang, 2015). Those too sure of their beliefs will be more vulnerable to other biases, such as naïve realism (Pronin, Gilovich, & Ross, 2004) or the “false consensus” effect (Krueger & Clement, 1994). Excessive faith in their beliefs can also lead people to discount others’ views (Minson & Mueller, 2012), or even disparage others as biased (Minson, Liberman, & Ross, 2011). Overprecision leads people to do too little to protect themselves against low-probability risks (Mannes & Moore, 2013). And overprecision blinds people to the need to consider other perspectives (Liberman, Minson, Bryan, & Ross, 2012; Ortoleva & Snowberg, 2015). These mistakes can have painful consequences for both individuals and organizations (Mergenthaler, Rajgopal, & Srinivasan, 2012). Given the costly consequences of overprecision, understanding its persistence is important.

We compare different measures of overprecision. Researchers have most often used the confidence interval paradigm employed by Alpert and Raiffa (1982). This method consistently finds overprecision (Bazerman & Moore, 2013), though it is controversial. The two most common critiques consider confidence intervals to be too difficult for participants to understand (Cosmides & Tooby, 1996), and that individuals do not naturally think about confidence in terms of confidence intervals (Mannes & Moore, 2013). We directly compare the confidence interval method with more naturalistic measures of confidence.

## Surprise

Surprise is one of the basic emotions (Levenson, 2011). Its fundamentally functional role is to highlight erroneous predictions and to direct attention at the surprising stimulus (Itti & Baldi, 2006). This function is sufficiently universal that surprise has proven useful in studying beliefs and expectations among monkeys and human infants (Steckenfinger & Ghazanfar, 2009; Xu & Spelke, 2000). When something unexpected happens, it receives more attention and longer gaze. The level of surprise one experiences and the duration of subsequent gaze is positively correlated with the degree of difficulty making sense of an event (Maguire, Maguire, & Keane, 2011). Seeing someone levitate is more surprising than seeing them jump. The more unexpected an event, the more intense the reaction to it (Mellers, Schwartz, Ho, & Ritov, 1997).

The more confident one is of one’s beliefs, the more surprising it should be when those beliefs turn out to be wrong. Given the ubiquity of overprecision in judgment, people should be surprised regularly. Here, we seek to connect the literature on surprise with the literature on overprecision in judgment. Research has not yet, to our knowledge, tested the hypothesis that predictions made with greater confidence will produce greater surprise when they turn out to be wrong. We test this prediction, and examine its consequences for the correction of subsequent confidence in judgment.

## Present Research

We asked participants to report ex-ante confidence for a variety of judgments, and after receiving performance feedback they then reported ex-post surprise at the result. This research hones in on a key interaction, wherein the relationship between ex-ante confidence and ex-post surprise is moderated by whether one’s answer is correct. We present three studies examining this relationship. Study 1 examines the relationships between confidence, correctness, and surprise for both self and others. Study 2 exogenously manipulates confidence and replicates the key finding from the first study. Study 2 uses a different manipulation of confidence and employs a repeated-measures design to explore the effect of confidence on surprise. Finally, in Study 3, we consider lay predictions regarding how surprised people believe they or others should be. Across the three studies, we employ a variety of different measures of belief precision and subsequent surprise, thereby testing their relationships with each other.

We report how we determined our sample size, all data exclusions (if any), all conditions, and all measures. Pre-registrations (for Studies 2 and 3), materials, and data are available: http://osf.io/j5vpe/.

## Study 1: Accuracy and Surprise

Study 1 examined the basic relationship between surprise and overconfidence. We hypothesized that both confidence and correctness would positively predict surprise. We also computed a measure of absolute distance from the true answer as an additional predictor on surprise. We expected higher ex-ante confidence would produce lower ex-post surprise for correct judgments, and for incorrect judgments higher ex-ante confidence would produce greater ex-post surprise. Study 1 seeks the antecedents of surprise, by examining the effect of confidence, correctness, and distance from the truth on subsequent surprise.

### Method

We exposed 430 MTurk workers to a set of five photographs of strangers. Participants had to guess how much each person weighed and report their confidence (on a scale from 0 – 100%) that their answer was within ten pounds of the true weight. After the five images, participants saw the true weight of each person in turn, followed by truthful feedback on whether or not their estimate fell within ten pounds of the actual weight. Following each round of feedback, participants reported their surprise on a scale of 0-100.

### Results and Discussion

Results reveal participants to be overprecise: The average hit rate for estimates within ten pounds of the true weight was 37%, yet average confidence was 66% (*SD* = 20.48). Given the repeated measures design, we employed a multilevel regression model with random slopes. We measured distance as the absolute value of the difference between the participants’ estimates and the true weight of the person in the photograph.

We conducted a multilevel regression predicting surprise from absolute distance, correctness, confidence, and the interaction between correctness and confidence, all nested at the individual level. The results reveal that absolute distance did indeed positively predict participants’ reported surprise, β = .33, *t*(451) = 14.84, *p* < .001. There were also significant main effects of confidence on surprise, wherein greater ex-ante confidence predicted greater ex-post surprise, β = .11, *t*(451) = 2.98, *p* = .003, and for correctness on surprise, β = .21, *t*(451) = 2.89, *p* = .003, where being correct predicted greater surprise. Additionally, there was a significant confidence-correctness interaction effect on surprise, β = −.21, *t*(451) = −2.82, *p* < .005.

We next compared correct and incorrect answers by analyzing them with separate multilevel regressions. When participants were incorrect, more confidence produced more surprise, β = .23, *t*(439) = 3.34, *p* < .001, and greater absolute distance from the truth also produced greater surprise, β = .52, *t*(439) = 10.47, *p* < .001. When participants were correct, confidence no longer predicted greater surprise, β = −.13, *t*(380) = −1.35, *p* = .17. However, the greater the distance between guesses and the truth, the more surprise they reported, β = .77, *t*(380) = 7.20, *p* < .001. See Figure 1.

**Figure 1.**
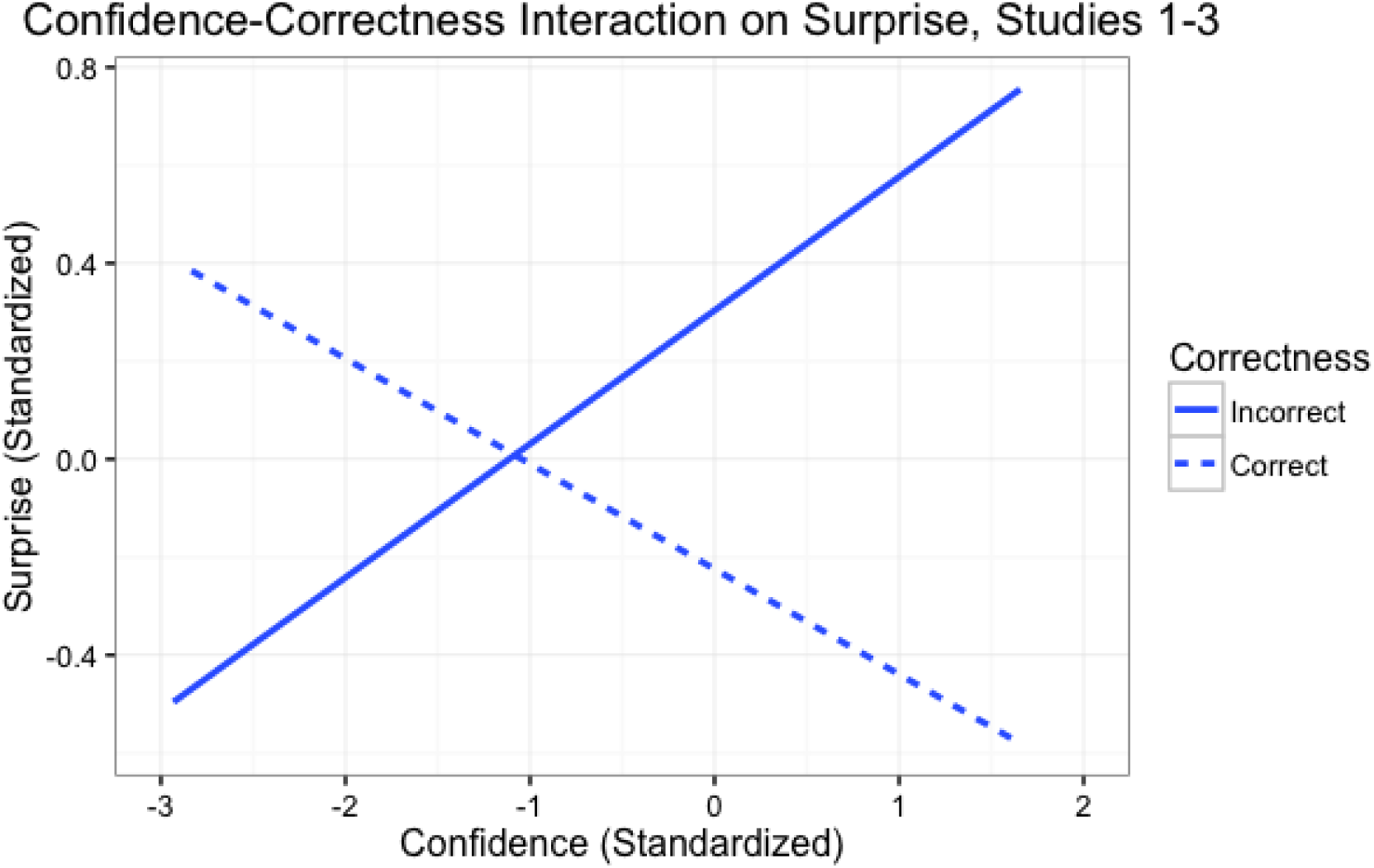
The aggregated standardized interaction effect of confidence and correct answers on reported ex-post surprise, for Studies 1-3. Measures are standardized by z-scoring within each study, then aggregating.

These results show that being confident and wrong is associated with greater surprise. However, this study is correlational in nature, leaving open the possibility that some third variable leads to both confidence and surprise. The next study employs an alternative measure of confidence and provides an experimental test of the effects of confidence on surprise using a longitudinal design.

## Study 2: Surprise Over Time

Study 2 sought to replicate Study 1’s results in a different domain while expanding our scope to two new facets of investigation. Specifically, we examined how aleatory or epistemic questions could affect the relationship between confidence and surprise. Because people are more confident when uncertainty is epistemic than when it is aleatory (Tannenbaum et al., 2016), this manipulation served as an experimental manipulation of confidence.

Ten questions were aleatory in nature, where the answer could not be known beforehand: What number ball would be drawn from the bingo cage? The other ten weight-guessing questions were epistemic in nature, with answers that participants could conceivably know, given sufficient skill at weight-guessing.

In addition, Study 2 sought to examine the temporal dynamics of confidence and surprise. Does experience help people adjust their expectations and better calibrate their confidence? Does it reduce surprise? We provided our participants with immediate feedback and measured the consequences of that feedback (reported surprise) on their subsequent reports of confidence.

### Method

Our pre-registered research plan called for 115 participants recruited via Amazon Mechanical Turk. We based this number on the effect size from a previous study (available in our <u>supplemental online materials</u>), f^2^ = .036, and power of 90%.

Study 2 asked each participant 20 questions; half aleatory and half epistemic. Our operationalization of aleatory questions came in the form of drawing bingo balls numbered from 1 to 100. Our epistemic questions asked participants to guess weights from photographs (as in Study 1). For each question, we asked participants to provide a best estimate of the answer, as well as high and low bounds to establish a confidence interval. Specifically, we asked for percentiles with only a 25% chance the true answer is above (or below) this high (or low) estimate. These interquartile ranges also defined which answers were considered ‘correct’ (i.e. whether the true answer was inside or outside of their self-created ‘confidence’ interval). Participants provided three responses per round: a best estimate plus high and low bounds, at the 25^th^ and 75^th^ percentiles, to form a 50% confidence interval. We used a standardized measure of the size of participants’ confidence intervals (their high estimate minus their low estimate divided by their ‘best guess’ estimate) as another confidence measure.

In addition, participants reported confidence on a 1 to 7 scale (from “*Not at all confident*” to “*Extremely confident*”). After learning the right answer, participants reported surprise on a 1-7 scale (from “*Not at all surprised*” to “*Extremely surprised*”).

### Results and Discussion

Hit rates between the 25^th^ and 75^th^ percentiles were similar for aleatory (65.83%) and epistemic questions (68.78%), *t*(114) = −1.51, *p* = .13. These hit rates suggest that participants were actually underprecise for both bingo ball questions and weight guessing questions, as both hit rates are greater than 50%. However, the bingo rounds produced lower feelings of confidence (*M* = 4.82, *SD* = 1.85) than did the weight rounds (*M* = 5.42, *SD* = 1.38), *t*(114) = −8.83, *p* < .001.

Using a standardized measure of confidence drawn from participants’ confidence intervals, dividing the range of their upper and lower estimates by their ‘best guess’ estimate, there appears a similar relationship: Participants provided larger confidence intervals on average for the bingo rounds (*M =* 64.85%, *SD* = 21.80%) compared to weight rounds (*M* =45.04%, *SD* = 19.29%), *t*(114) = 23.07, *p* < .001.^1^ There was also a difference in reported surprise by question type, *t*(114) = −8.10, *p* < .001. Participants reported greater surprise for epistemic weight-guessing judgments (*M* = 3.40, *SD* = 1.91) than for aleatory bingo-ball judgments (*M* = 2.71, *SD* = 2.15), *t*(114) = 8.10, *p* < .001.

The effect of confidence and correctness on surprise replicates the results of Study 1: when incorrect, high confidence predicted greater ex-post surprise, *β* = .33, *t*(114) = 9.37, *p* < .001; though when correct, higher confidence predicted lower ex-post surprise, β = −.33, *t*(114) = −12.89, *p* < .001.

A multilevel regression predicting surprise with question type, correctness, distance, confidence, and a correctness-confidence interaction reveals a significant main effect of question type: the epistemic weight-guessing questions elicited greater surprise than the aleatory bingo ball questions, β = .15, *t*(119) = 9.63, *p* < .001. This analysis also replicated the confidence-correctness interaction from previous studies, β = −1.33, *t*(119) = −23.52, *p* < .001. This interaction arises because confidence increased surprise only following incorrect answers. Subsetting on correctness identifies the negative relationship between confidence and surprise when correct, β = −.33, *t*(117) = −12.89, *p* < .001, and the positive relationship between confidence and surprise when incorrect, β = .33, *t*(115) = 9.37, *p* < .001. See Figure 1.

However, our other measure of confidence (confidence interval size) produces very different results. The same multilevel model predicting surprise with question type, correctness, distance, and confidence (from interval size), produces a correctness-confidence interaction, β = .31, *t*(119) = 5.16, *p* < .001. Again to seek clearer insights, we ran this new model on data subsetted by correctness. The negative relationship between confidence and surprise for correct answers is weakened to nonsignificance, β = −.04, *t*(119) = −1.55, *p* = .12. The positive relationship we found previously between confidence and surprise when incorrect is nearly nonexistent, β = .02, *t*(119) = 0.59, *p* = .56.

The relationship between confidence interval size and self-reported confidence is not intuitive for participants. Confidence interval size was correlated with self-reported confidence at *r* = .28, *p* < .001. This suggests that as intervals grew (implying lower confidence), self-reported confidence also increased--a contradictory pattern. Puzzlingly, this correlation between CI size and scale confidence only exists with incorrect answers, *r* = .30, *p* < .001, the correlation is nonsignificant for correct answers, *r* = .05, n.s. This is intriguing as both measures of confidence are reported prior to participants learning the correct answer.

This study’s repeated-measures design allowed us to test the effect of surprise on confidence over time. We employed a lagged regression to see how confidence is predicted by correctness at *t* – 1 (i.e., how correctness in any given round predicts confidence in the round immediately following). Lagged correctness at *t* – 1 significantly predicted confidence at time *t*, β = .07, *t*(115) = 4.12, p < .001. In other words, participants expanded their confidence intervals after having been wrong. Did reported surprise impact subsequent confidence similar to how correctness affected subsequent confidence? Adding lagged surprise at *t* – 1 into the same model showed no significant effect of lagged surprise on confidence, β = .003, *t*(116) = 0.13, *p* = .89. This result suggests a profound failure of the functional role of surprise: it did not reduce subsequent confidence. The inclusion of lagged surprise also left the main effect of lagged correctness on surprise nearly unchanged, β = .07, *t*(116) = 3.78, *p* < .001.

Results of Study 2 introduce important caveats to our key interaction between confidence and correctness on reported surprise. The results reinforce the importance of the method one uses to elicit confidence, as our key results largely disappear using the confidence-interval elicitation. This might be attributable to the difficulties people have setting confidence intervals. For instance, people set 50% confidence about as wide as they set 98% confidence intervals, despite the fact that 98% confidence intervals should be much wider (Teigen & Jørgensen, 2005). These results add to our skepticism of confidence-interval measures and the degree to which they effectively capture subjective feelings of confidence. However, there is another possibility: Because two effects may have cancelled each other out: (1) more confident people were more surprised at being wrong but (2) people who gave wider intervals (i.e. those less confident) were more surprised that their larger intervals did *not* contain the right answer. This ambiguity interpreting confidence interval measures led us to abandon confidence interval elicitations in Study 3.

## Study 3: Predicting Surprise

Study 3 asks whether people are as surprised as they *should* be. The study randomly assigned half the participants to a prediction condition which elicited predictions of surprise for hypothetical correct and incorrect answers. The other half of participants, in the control condition, made no such predictions. Study 3 again employed both aleatory and epistemic judgments.

In choosing which tasks to use, we noted that Study 2 found surprise to be lower for bingo balls than weight-guessing. Since the distribution of bingo balls is uniform, all numbers are equally likely and there is little reason to be surprised by any particular outcome, potentially contributing to diminished “surprisingness.” Therefore, Study 3 replaces bingo ball draws with a set of ten coin flips, which has a single-peaked distribution of outcomes and the potential for truly surprising outcomes (such as ten flips all coming up tails).

### Method

A power analysis of the prior studies, where people reported ex ante surprise providing average effect sizes of f^2^=.036 to f^2^=.024 provided a recommended sample size of 105. Wary of losing power from subsetting the data in testing our hypotheses, we pre-registered a sample size of 150 and ended up with 151 participants.

Each survey informed participants that there would be ten rounds of ten coin flips each and ten rounds of guessing individuals’ weights from images. For each of the two blocks (which were presented in a random order) participants provided a best guess estimate as well as their confidence in being correct (on a 1-100 scale). Following each round, we truthfully informed participants whether they had answered correctly, and then participants reported their surprise at the outcome (on a 1-7 scale). Answers for coin flips counted as correct if they were within one head (out of ten) of the actual outcome. Weight guesses counted as correct if they were within ten pounds of the truth.

We assigned participants to one of two between-subjects conditions: a control condition and a prediction condition, where, in each round, in addition to the procedure described above, participants predicted how surprised they would be if their answer was right and if it was wrong.

### Results and Discussion

On average, participants were overconfident. They report being 55.5% confident on average, but they are only right 50.3% of the time, one-sample *t*(2804) = 12.1, *p* < .001. This confidence declines with experience. It starts at 60.2% in Round 1 and declines to 52.8% in Round 20. A linear regression predicting confidence with round number, coin/weight, and fixed effects for subject produces a strong effect of round, *B* = −.59, *t* = −5.08, *p* < .001. There is no significant difference in expressed confidence between weight and coin rounds, *B* = −1.55, *t* = −1.16, *p* = .25. This lack of a difference between weight and coin rounds is remarkable given that participants’ guesses are correct 61% of the time for coin rounds but only 39.8% of the time for weight rounds, *t*(3038) = 11.83, *p* < .001.

Were participants as surprised as they predicted they should be? In order to test this, we employed two independent-samples t-tests to account for the repeated measures design, comparing the predicted surprise for a correct answer to a subsequent correct answer’s surprise, and comparing the predicted surprise for an incorrect answer to a subsequent incorrect answer’s surprise. Through this we see that participants reported less surprise (*M* = 3.30, *SD* = 1.91) than they predicted they would (*M* = 3.78, *SD* = 1.82) when correct, *t*(712) = 7.63, *p* < .001. Conversely, they were more surprised *(M* = 4.77, *SD* = 1.82) than predicted (*M* = 3.61, *SD* = 1.74) when incorrect, *t*(761) = −15.56, *p* < .001. These patterns are similar for both coin and weight rounds.

We also explored whether the mere act of prediction had an impact on one’s subsequent surprise. We employed a repeated measures one-way ANOVA predicting ex-post surprise by condition, controlling for individual-level error. This analysis shows that condition significantly predicts ex-post surprise, *F*(1, 2888) = 19.41, *p* < .001. A follow up hierarchical linear regression controlling for individual-level error shows that being in the prediction condition increased ex-post reported surprise, β = 0.17, *t*(150) = 4.41, *p* < .001. Those who predicted their surprise wound up reporting more surprise (*M* = 4.04, *SD* = 2.01) than those who did not (*M* = 3.39, *SD* = 2.04).

In testing a replication of the lagged analysis from Study 2 we ran a similar lagged analysis predicting confidence with correctness at *t* – 1. The analysis shows that when it was the only predictor, lagged correctness positively predicted subsequent confidence, β = .04, *t*(150) = 2.81, *p* < .005. However, including lagged surprise wipes out the relationship, leaving only lagged surprise significant, β = −0.09, *t*(150) = −4.23, *p* < .001, where more surprise at *t*-1 led to less confidence at *t.* This contrasts with the result from in Study 2.

To explore whether this failure to replicate was an artifact of eliciting predictions of surprise in the prediction condition, we subsetted the data by condition and ran the same analyses. In the prediction condition, the relationship stayed significant with less power, β = −.08, *t*(75) = −2.70, *p* = .007. The relationship also holds in the control condition, which presents itself as a close replication of Study 2 since participants made no predictions, β = −.10, *t*(73) = −3.45, *p* < .001. In short, results from Study 3 do indeed document the corrective effect by which surprise reduces subsequent confidence.

## GENERAL DISCUSSION

Our results show that ex-ante confidence and ex-post surprise are inextricably linked. Our primary finding is that when people are correct, greater ex-ante confidence produces less ex-post surprise, whereas when they are incorrect, greater ex-ante confidence produces more ex-post surprise. We examine the psychology underlying these relationships and identify moderators that can either suppress or enhance their strength. Studies 1 and 2 establish the link between confidence and surprise, highlighting that correctness is a powerful moderator of the relationship. Studies 2 and 3 employ exogenous manipulations of confidence; their results replicate the correlational results of Study 1. Study 2 finds more powerful confidence-correctness interaction effects on surprise for epistemic questions than for aleatory, consistent with the notion that feeling personally accountable for knowing or not knowing the answer increases the intensity of emotional reactions to being right or wrong. Study 3 finds that people are more surprised about being wrong than they expect to be.

What of the utility of surprise? If surprise reflects prediction error, individuals should seek to maximize accuracy and minimize surprise (Ely, Frankel, & Kamenica, 2015). This implies that surprise should lead people to reduce their subsequent confidence. Our results suggest that surprise does not always play this functional role, or that it is difficult to measure consistently. Future research should examine the conditions under which surprise has a corrective effect on subsequent confidence. How quickly does this effect decay and what possible moderators could increase the calibrating power and longevity of feedback on subsequent confidence? Could incorrect answers in epistemic domains more central to one’s self-concept ‘stick’ for a longer period of time, forcing one’s re-evaluation of their believed expertise? Or could the opposite be the case, where the incorrect answer is considered anomalous and the sense of expertise persists?

We aspired to measure the effects of overprecision on surprise. In recording participants’ ex-ante confidence, their correctness, and their ex-post surprise, we document consistent evidence suggesting that people expect to be correct. If they go into a decision with confidence, they are more surprised to be incorrect, and less surprised when correct. We believe these results do more than underscore precision in judgment. Rather, this research approaches the topic with a new paradigm that serves to reveal another layer in the scientific understanding of the psychology of confidence and precision in judgment.

## Footnotes

1 This result follows from the fact that the range for bingo balls runs from 0 to 100, whereas the functional range for weights runs from 100 to 200. Consequently, intervals of similar size are larger in relation to best guesses for bingo balls. So the result probably does not qualify as psychologically meaningful.

